## Supplementary figures and images for "Assessment of the Histone Mark-based Epigenomic Landscape in Human Myometrium at Term Pregnancy"

### Supplementary Figure 1

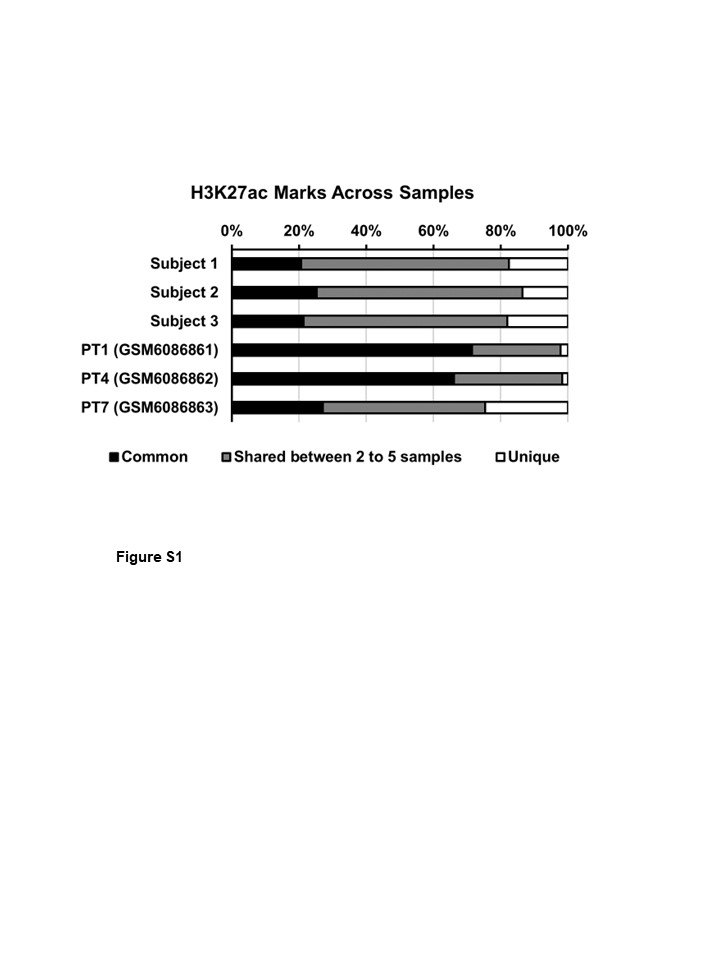

### Supplementary Figure 2

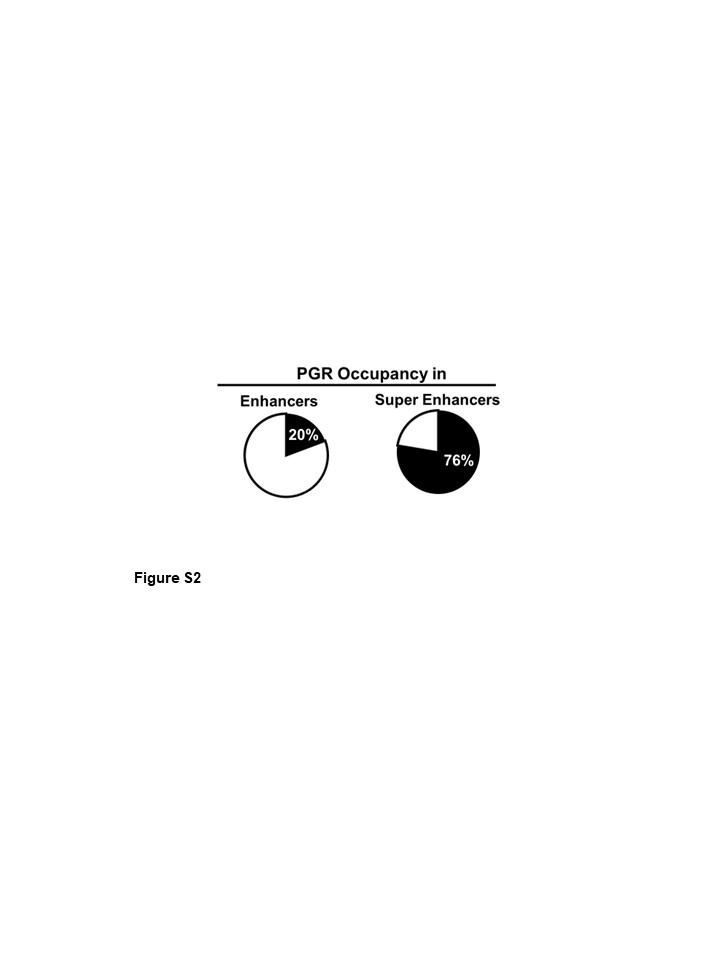
